# 4D intravital imaging studies identify platelets as the predominant cellular procoagulant surface in a mouse model of hemostasis

**DOI:** 10.1101/2023.08.25.554449

**Authors:** Abigail Ballard-Kordeliski, Robert H. Lee, Ellen C. O’Shaughnessy, Paul Y. Kim, Summer Jones, Nigel Mackman, Matthew J. Flick, David S. Paul, David Adalsteinsson, Wolfgang Bergmeier

**Affiliations:** Department of Biochemistry and Biophysics, University of North Carolina at Chapel Hill, NC, USA; UNC Blood Research Center, University of North Carolina at Chapel Hill, NC, USA; Department of Cell Biology and Physiology, University of North Carolina at Chapel Hill, NC, USA; Department of Medicine, McMaster University, Hamilton, ON, Canada; Thrombosis and Atherosclerosis Research Institute, Hamilton, ON, Canada; Department of Medicine, University of North Carolina at Chapel Hill, NC, USA; Department of Pathology and Laboratory Medicine, University of North Carolina at Chapel Hill, NC, USA; Department of Mathematics, University of North Carolina at Chapel Hill, NC, USA

## Abstract

Interplay between platelets, coagulation/fibrinolytic factors, and endothelial cells (ECs) is necessary for effective hemostatic plug formation. This study describes a novel four-dimensional (4D) imaging platform to visualize and quantify hemostatic plug components with high spatiotemporal resolution. Fibrin accumulation following laser-induced endothelial ablation was observed at the EC-platelet plug interface, controlled by the antagonistic balance between fibrin generation and breakdown. Phosphatidylserine (PS) was first detected in close physical proximity to the fibrin ring, followed by exposure across the endothelium. Impaired PS exposure in *cyclophilinD*^*-/-*^ mice resulted in a significant reduction in fibrin accumulation. Adoptive transfer and inhibitor studies demonstrated a key role for platelets, but not ECs, in fibrin generation during hemostatic plug formation. Inhibition of fibrinolysis with tranexamic acid (TXA) led to increased fibrin accumulation in WT mice, but not in *cyclophilinD*^*-/-*^ mice or WT mice treated with antiplatelet drugs. These studies implicate platelets as the functionally dominant procoagulant surface during hemostatic plug formation. In addition, they suggest that impaired fibrin formation due to reduced platelet procoagulant activity is not reversed by TXA treatment.

## INTRODUCTION

Hemostatic plug formation is critical for the prevention of blood loss from sites of vascular injury. Platelets are rapidly recruited to the site of injury and form a loose platelet plug. Consolidation of the platelet plug depends on coagulation activation, thrombin generation, and the formation of a fibrin matrix (1). Fibrin degradation and plug resolution are dependent on the recruitment of plasminogen and the generation of plasmin, the key enzyme of the fibrinolytic system. Both platelets and endothelial cells (ECs) are known modulators of fibrin formation and fibrin breakdown, and both thrombin and plasmin have documented roles in platelet and EC activation (2–4). Most of this knowledge is derived from in vitro studies with purified platelets and cultured ECs or ex vivo studies under flow conditions, utilizing microfluidic-based thrombus formation assays (5,6).

Upon strong stimulation, platelets provide a negatively charged surface for the binding of coagulation factors, especially the positively charged GLA domain-containing proteins prothrombin, FVII, FIX, and FX. Critical for the interaction of these coagulation factors with the cell surface is the translocation of negatively charged phospholipids, such as phosphatidylserine (PS), from the inner to the outer leaflet of the plasma membrane (PM), facilitated by various cellular changes, including the depolarization of the mitochondrial membrane and prolonged high cytosolic calcium levels (7,8). Cyclophilin D (CypD) is an important regulator of the mitochondrial permeability transition pore (MPTP), and platelets lacking CypD are markedly impaired in their ability to express PS on their surface (9). Elevated calcium, facilitated by mitochondrial depolarization, triggers the activation of the phospholipid scramblase, transmembrane protein 16F (TMEM16F), which facilitates PS translocation to the surface (10,11). TMEM16F was identified as the gene affected in Scott Syndrome, a rare congenital hemorrhagic disorder (12). Studies in animal models have demonstrated a key role for both platelet and EC TMEM16F in thrombus formation (13,14).

ECs also support prothrombinase assembly and thrombin generation in vitro (15,16). Importantly, several recent intravital microscopy imaging studies have suggested that ECs play a major role in thrombin generation at sites of vascular injury. Ivanciu et al. reported a prominent contribution of the activated endothelium, but not platelets, to prothrombinase complex assembly and fibrin formation adjacent to the site of laser damage in cremaster arterioles (17). More recently, Schmaier et al. reported vessel wall-dependent fibrin formation on injured cremaster arterioles, with both TMEM16F and TMEM16E contributing to the procoagulant activity and thrombus formation at the site of injury (13). However, both studies used indirect means to reduce/limit platelet procoagulant activity, and vascular injuries were minor, not requiring plug formation to secure hemostasis.

Once formed, tissue plasminogen activator (tPA) and plasminogen bind to and degrade fibrin in the plug. Studies in vitro and in vivo identified a key role for platelets in fibrinolysis (18,19). Platelets were found to affect both the quantity and the quality of the fibrin network, with fibers around platelets being thin, densely packed, and more resistant to lysis compared to fibers that were not associated with platelets (20). In vitro and in vivo studies also suggest that plasminogen recruitment to platelets is almost entirely dependent on fibrin (19,21). Platelets also express the plasminogen receptor, Plg-RKT, and in vitro studies suggest a significant role for this receptor in plasminogen retention on the activated platelet membrane (22). Like platelets, ECs were shown to both positively and negatively regulate fibrinolysis (4). ECs constitutively release tPA at low concentrations, which is retained on their surface and plays a role in promoting fibrinolysis in a fibrin-dependent manner (23). Urokinase-type plasminogen activator is also released by ECs upon stimulation. Interestingly, ECs also express and release plasminogen activator inhibitor 1 (PAI-1) and thus can downregulate fibrinolysis under certain conditions (24). These data suggest that selective up- or down-regulation of these factors would dictate the net fibrinolytic activity surrounding the ECs.

To date, no study has examined on a molecular level, in vivo, how platelets, plasma proteins and ECs interact during hemostatic plug formation. To do so, we established a novel 4-dimensional (4D) imaging platform, which we then used to visualize platelets, ECs, PS, fibrin, and plasminogen during hemostatic plug formation with high spatiotemporal resolution. Using this new imaging modality in combination with knockout/inhibitor-treated mice and an adoptive platelet transfer method (25), we demonstrate that 1) PS is first expressed on adherent platelets, coinciding in space and time with the accumulation of fibrin, 2) PS, but not fibrin, then is detected across the endothelium near the injury site, 3) fibrin accumulation is markedly reduced in *CypD*^*-/-*^ mice and mice lacking CypD in platelets only, 4) recombinant plasminogen predominantly colocalizes with fibrin at the injury site, 5) increased fibrin accumulation on the EC surface is observed in plasminogen knockout (*Plg*^*-/-*^) mice, and 6) tranexamic acid (TXA) leads to increased fibrin accumulation in WT mice, but not in WT mice with impaired platelet procoagulant response.

## RESULTS

### An In Vivo 4D Imaging Model of Hemostasis

The saphenous vein laser injury model is a well-established method for imaging hemostasis in mice (26). Upon exposure of the saphenous vein (Figure 1A), laser ablation is used to create a penetrating vascular lesion with a diameter of ∼50 μm (Figure 1B). Traditionally, this model was limited to three dimensions, either 2D over time or 3D at a fixed time point. We here utilized time-lapse spinning disk confocal microscopy to quantify enrichment of plug components with improved spatiotemporal resolution. Injuries analyzed in this study ranged from 40-60 μm in diameter. Images for a 150 μm Z-stack (Figure 1C) were sequentially recorded at a step size of 7.5 μm, requiring 4.33 seconds per fluorescence channel. Post-acquisition generation and analysis of 4D movies was performed in Imagetank software (27). Mice with GFP-expressing platelets and tdTomato-expressing endothelium (mTmG x PF4-Cre mice) were injected with AlexaFluor(AF)647-labeled anti-fibrin antibodies and hemostatic plug components were visualized following laser injury. Fibrin accumulation was rapidly observed at the perimeter of the platelet plug, forming a ring at the level of the endothelium (Figure 1D; supplemental video 1). While the intensity of the fibrin signal increased throughout the observation period, the ring pattern did not change. The localization of PS was also examined in mTmG x PF4-Cre mice injected with AF647-labeled Annexin-V (aV). At the 1-minute time point after injury, PS exposure was largely localized to the edge of the platelet plug, similar to where we observed the fibrin ring (Figure 1E; supplemental video 2). At later time points, however, PS was detected across both the intraluminal and the extraluminal surface of the endothelium. The difference in fibrin versus PS signal in relation to the endothelium and the platelet plug is illustrated in Figure 1F, which shows overlays of histograms for each fluorescence channel at t = 1, 5, and 13 min after injury.

**Figure 1.**
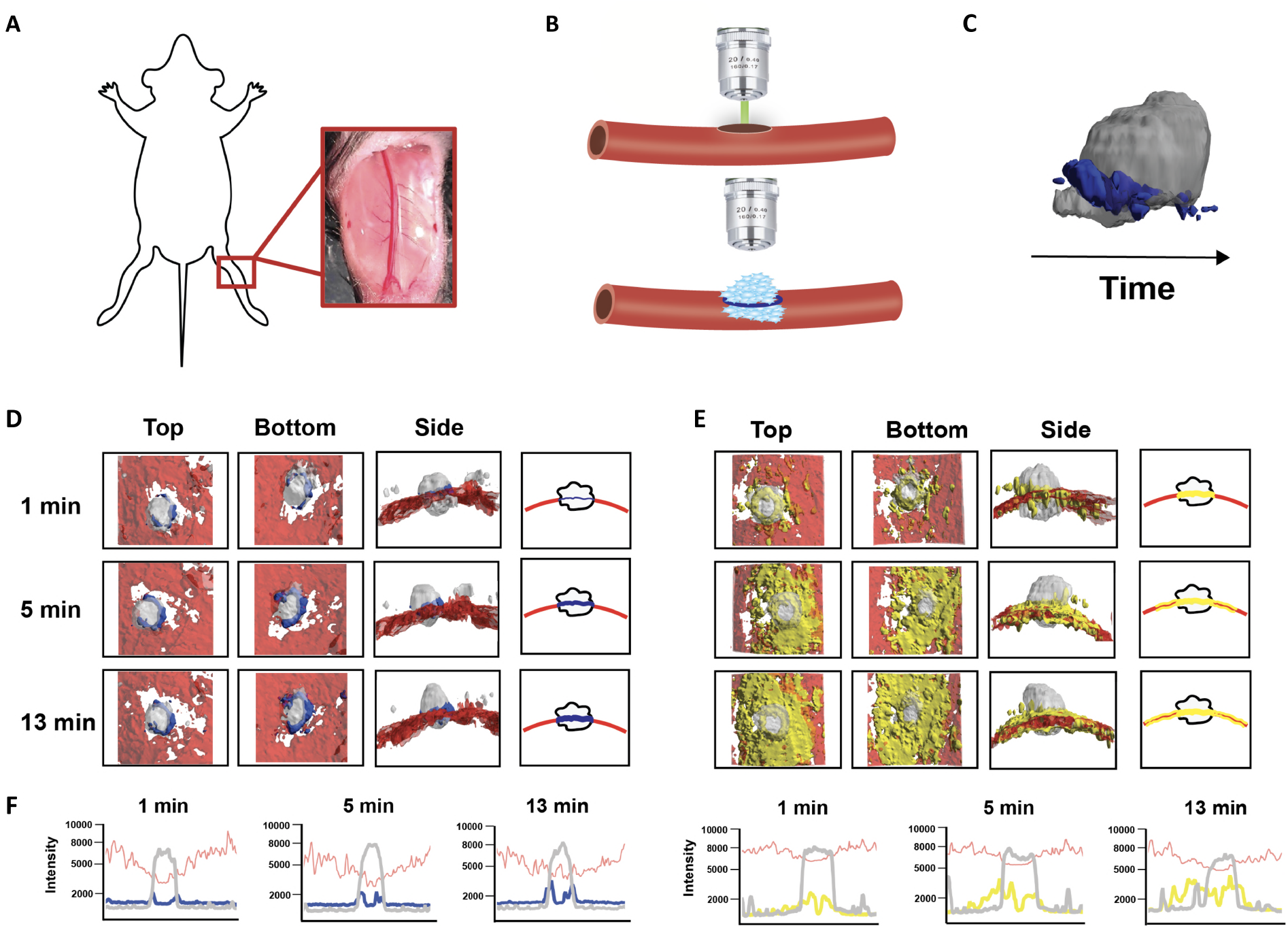
4D imaging model of hemostasis. **(A-C)** Experimental set-up. **(A)** Exposed saphenous vein. **(B)** Laser-induced endothelial ablation of saphenous vein was induced with Ablate! photoablation system with a 532-nm pulse laser. Accumulation and localization of various hemostatic plug components were recorded using a Zeiss Axio Examiner Z1 microscope with a 20x/1 numerical aperture water immersion objective lens and Slidebook software. **(C)** Sequential 150 μm Z-stacks of the forming hemostatic plug were acquired with 7.5 μm steps and 4 x 4 binning. Image analysis was performed with Image Tank Software (Visual Data Tools). Image shows surface view of platelets (grey) labeled with anti-GPIX-AlexaFluor(AF)488 and fibrin (blue) labeled with anti-fibrin-AF647 antibodies. **(D and E)** Visualization of platelets, endothelial cells, fibrin, and phosphatidylserine (PS) within forming hemostatic plugs at the indicated time points following laser injury in cell lineage tracer mice with green fluorescent protein (GFP)-expressing platelets and tdTomato-expressing endothelium. Views shown: Top = top down (extravascular) view; Bottom = bottom up (intravascular) view; Side = side view; right panel: schematic representation for side view images. **(D)** Localization of fibrin (blue) in hemostatic plugs of mice injected with anti-fibrin-AF647 antibody. **(E)** Localization of PS-positive surfaces (yellow) at sites of injury in mice injected with AnnexinV-AF647. **(F)** Overlay of fluorescence intensity histograms for platelets (grey), ECs (red) and fibrin (blue, left panel) or PS (yellow, right panel), measured along one cross-sectional slice of a Z-max projection.

### Plug fibrin accumulation is determined by an antagonistic balance between fibrin formation and fibrin degradation

To validate that this new model faithfully reports thrombin/ fibrin generation at the site of hemostatic plug formation, we next studied mice expressing a low level of tissue factor (TF) (*TF*^*low*^) (28) and mice deficient in coagulation factor XII (*F12*^*-/-*^) (29). Consistent with the key role of TF in coagulation initiation, we observed a markedly delayed and significantly reduced accumulation of fibrin within hemostatic plugs in *TF*^*low*^ mice (Figure 2A,B; supplemental video 3). Similar to controls, the limited fibrin accumulation still occurred at the edge of the platelet plug. Platelet accumulation was not reduced (Figure 2C), while platelet density in these smaller plugs was slightly reduced when compared to control plugs (Figure 2D). Fibrin accumulation was also significantly reduced in *F12*^*-/-*^ mice (Figure 2E,F; supplemental video 4). The initial rise in fibrin levels was reduced but not abolished, suggesting that the first wave of TF/FVIIa-mediated thrombin generation remains largely intact in *F12*^*-/-*^ mice. Overall platelet accumulation in *F12*^*-/-*^ plugs was not altered (Figure 2G), while plug density was significantly lower than in controls (Figure 2H). Together, these findings are consistent with a key role for the extrinsic pathway in the initiation of thrombin generation following laser injury, while FXII is critical for subsequent contact pathway activation.

**Figure 2.**
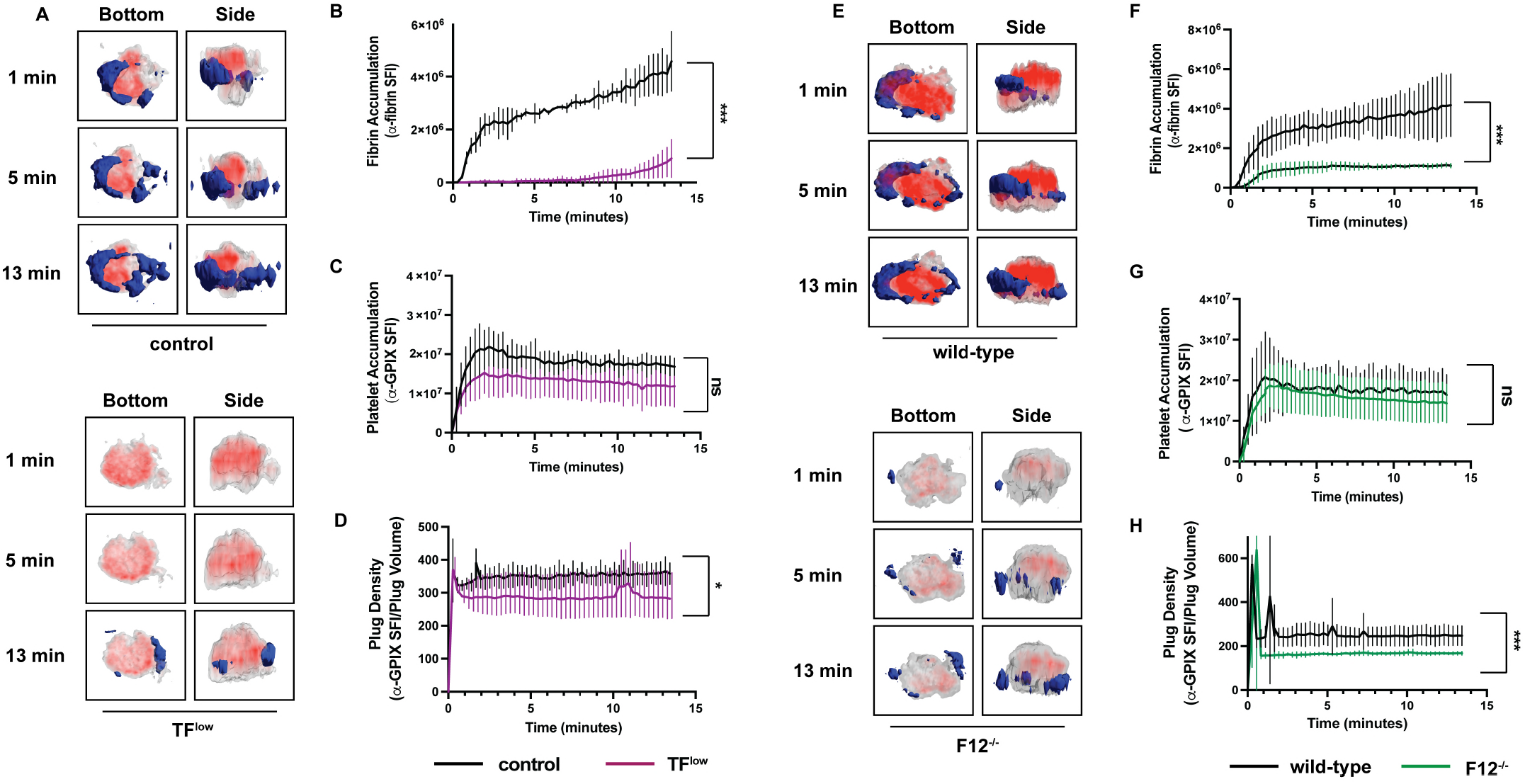
Both tissue factor (TF) and FXII are critical for fibrin accumulation. (A) Platelet (anti-GPIX-AF488) and fibrin (anti-fibrin-AF647) accumulation at injury sites in control (top panel) and *TF*^*low*^ mice (bottom panel). Representative images showing surface view for fibrin ring (blue) around platelet plug (grey), overlayed with platelet density view (red). **(B)** Sum fluorescence intensity (SFI) ± standard error of the mean (SEM) for fibrin signal. (*TF*^*low*^ n=16, control n=14) **(C)** SFI ± SEM for platelet signal. (*TF*^*low*^ n=16, control n=12) **(D)** Plug density as determined by the ratio between anti-GPIX SFI and plug volume. (*TF*^*low*^ n=16, control n=12) **(E)** Platelet (anti-GPIX-AF488) and fibrin (anti-fibrin-AF647) accumulation at injury sites in *F12*^*+/+*^ controls (top panel) and *F12*^*-/-*^ mice (bottom panel). Representative images shown as in (A). **(F)** SFI ± SEM for fibrin signal. (*F12*^*-/-*^ n=19, *F12*^*+/+*^ =23) **(G)** SFI ± SEM for platelet signal. (*F12*^*-/-*^ n=17, *F12*^*+/+*^ =19) **(H)** Plug density determined as in (D). (*F12*^*-/-*^ n=17, *F12*^*+/+*^ =19) \**P* < .05,\*\*\**P* < .001. ns, not significant.

We next studied how deficiency in plasminogen (*Plg*^*-/-*^ mice) (30) affects fibrin accumulation in hemostatic plugs. As expected, decreased fibrinolysis in *Plg*^*-/-*^ mice resulted in significantly increased fibrin accumulation within plugs (Figures 3A,B; supplemental video 5). Interestingly, the increase in fibrin accumulation was observed at all time points throughout the observation period. The additional fibrin was observed at the extravascular interface of the plug and intravascularly along the endothelial surface (Figures 3A). Consistently, both increased fibrin surface area (Figure 3C) and increased fibrin ring thickness (Figure 3D) were observed in *Plg*^*-/-*^ mice. Intensity plots overlaying the fluorescence intensity for platelets and fibrin were created to further illustrate the expansion of fibrin accumulation onto the endothelium, away from the platelet plug (Figure 3E; supplemental video 6).

**Figure 3.**
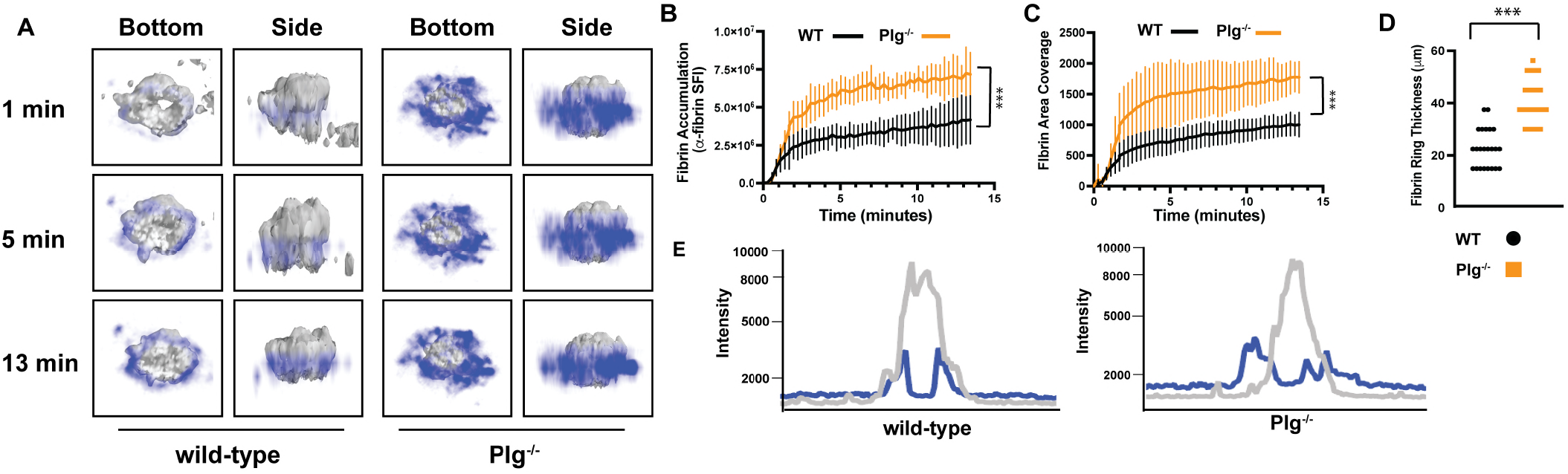
Increased fibrin accumulation in plasminogen-deficient mice. **(A)** Representative images for fibrin (blue, intensity view) and platelet (grey, surface view) accumulation at the indicated time points after laser injury in *Plg*^*+/+*^ controls (left panel) and *Plg*^*-/-*^ mice (right panel). **(B)** SFI ± SEM for fibrin signal. (Plg^+/+^ n=12, *Plg*^*-/-*^ n=13) **(C)** Fibrin area coverage. **(D)** Fibrin ring thickness (μm) measured at t=13 minutes after laser injury. **(E)** Overlay of fluorescence intensity histograms for platelets (grey) and fibrin (blue) in plugs from a *Plg*^*+/+*^ (left panel) or *Plg*^*-/-*^ mouse (right panel), measured along one cross-section slice of a Z-max projection. \*\*\**P* < .001. ns, not significant.

### Platelet Mediated Procoagulant Response is Critical for Fibrin Accumulation

CypD, an important regulator of the MPTP opening and subsequent PS surface exposure, is expressed in platelets and ECs (9,31). To study the impact of impaired PS exposure on hemostatic plug formation, we analyzed wild-type (WT) and CypD knockout (*CypD*^*-/-*^) mice in the saphenous vein laser injury model with 4D imaging. As shown in Figure 1, aV staining and fibrin accumulation co-localized one minute after laser injury to the saphenous vein of WT mice, and then the aV signal continued to spread across the endothelium while fibrin remained constrained to the ring (Figure 4A; supplemental video 7). *CypD*^*-/-*^ mice exhibited markedly decreased PS exposure with no PS near the platelet plug at early time points after injury and minimal PS on the endothelium at later time points (Figures 4A,C). Consistent with a key role of PS-positive surfaces in the assembly and activation of coagulation factors, fibrin accumulation at sites of laser injury leveled off early and was significantly reduced in *CypD*^*-/-*^ mice (Figures 4B,D). Platelet accumulation was not significantly different between *CypD*^*-/-*^ and control mice (Figure 4E).

**Figure 4.**
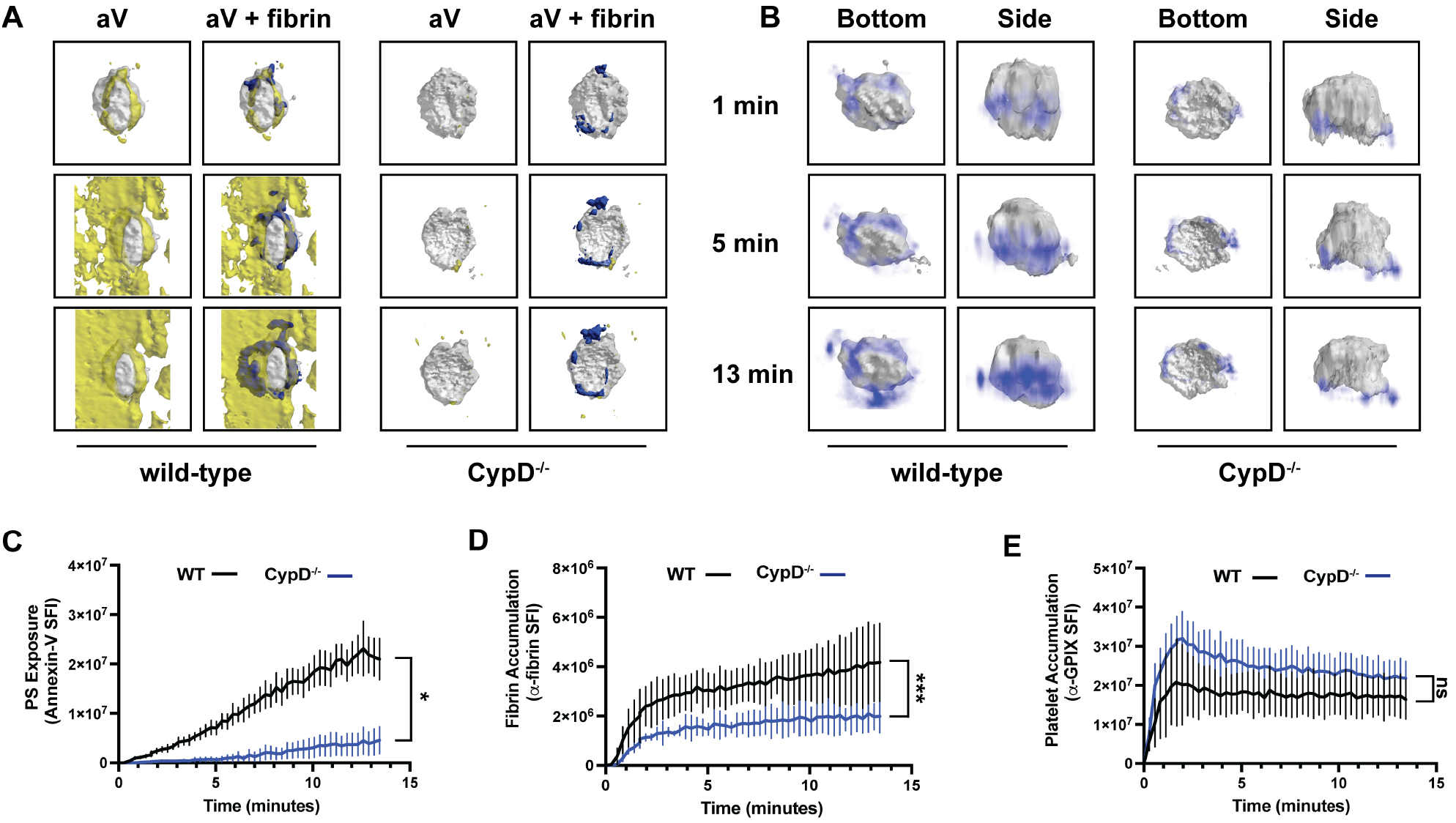
PS exposure potentiates fibrin accumulation around the platelet plug. **(A)** Surface view for accumulation of fibrin (blue), PS (yellow), and platelets (grey) at the indicated time points during hemostatic plug formation in wild-type (left panel) and *CypD*^*-/-*^ (right panel) mice. PS was labeled with AF568-conjugated aV. **(B)** Fibrin (blue, intensity view) and platelet (grey, surface view) accumulation at the indicated time points during hemostatic plug formation in wild-type (left panel) and *CypD*^*-/-*^ (right panel) mice. **(C-E)** SFI ± SEM over time for **(C)** PS (wt n=7, *CypD*^*-/-*^ n=8), **(D)** fibrin (wt n=20, *CypD*^*-/-*^ n=25), and **(E)** platelets (wt n=20, *CypD*^*-/-*^ n=27). \*\**P* < .01; \*\*\**P* < .001. ns, not significant.

To differentiate the relative roles of platelet versus EC PS in fibrin accumulation at sites of vascular injury, we next performed studies in mice lacking CypD in platelets only. These mice and appropriate controls were generated by adoptive transfer of *CypD*^*-/-*^ or WT platelets into thrombocytopenic mice with normal CypD expression in ECs (25,32). The spatiotemporal characteristics of aV staining and fibrin accumulation at sites of laser injury were comparable between thrombocytopenic mice receiving WT platelets (both EC and platelets express CypD) (Figure 5A-D; supplemental Video 8) and non-depleted, non-transfused WT mice (Figure 4). A lack of CypD in platelets only did not significantly impair total PS expression in the vicinity of the hemostatic plug (Figures 5A,C). However, a marked reduction in platelet-associated PS was observed at early time points after laser injury (see “1 min” images in Figure 5A; supplemental Video 9). Significantly reduced fibrin accumulation (Figures 5B,D) but normal platelet accumulation (Figure 5E) were observed in mice lacking CypD in platelets only, similar to what we observed in *CypD*^*-/-*^ mice (Figure 4). These data indicate that platelet PS, and not EC PS, is the main contributor to thrombin generation and fibrin accumulation in hemostatic plugs.

**Figure 5.**
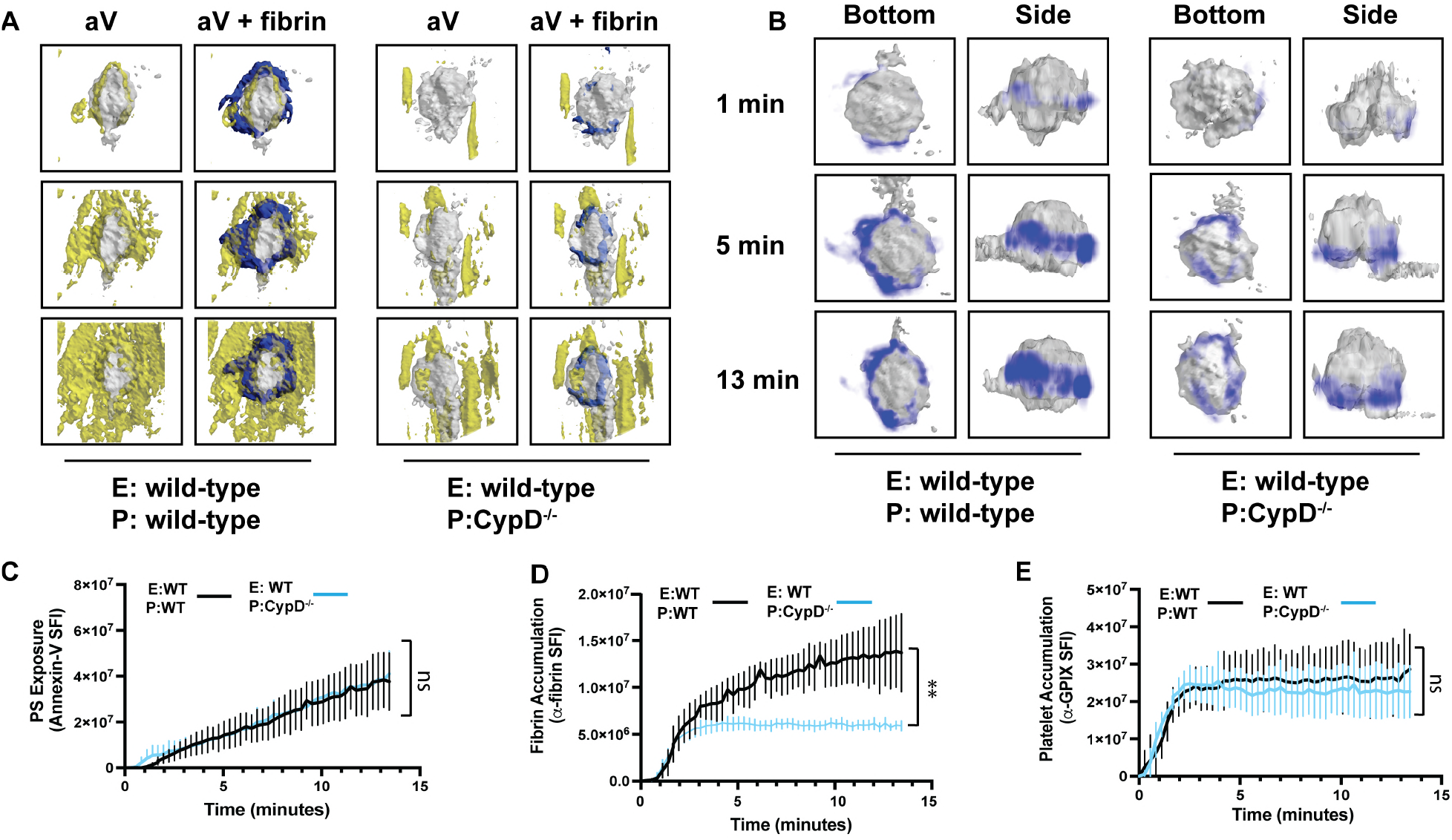
Platelet PS exposure is critical to fibrin accumulation during hemostatic plug formation. Mice lacking CypD in platelets only were generated by adoptive transfer of *CypD*^*-/-*^ platelets into thrombocytopenic recipient mice (see Methods). **(A)** Surface view for accumulation of fibrin (blue), PS (A5, yellow), and platelets (grey) at the indicated time points during hemostatic plug formation in E:wt/P:wt (left panel) and E:wt/P:CypD^-/-^ (right panel) mice. **(B)** Fibrin (blue, intensity view) and platelet (grey, surface view) accumulation at the indicated time points during hemostatic plug formation in E:wt/P:wt (left panel) and E:wt/P:CypD^-/-^ (right panel) mice. **(C-E)** SFI ± SEM over time for **(C)** PS (E:wt/ P:wt =5, E:wt/P:CypD^-/-^ =5), **(D)** fibrin (E:wt/P:wt: n =15; E:wt/P:CypD^-/-^; n=17), and **(E)** platelets (E:wt/P:wt n=12; E:wt/ P:CypD^-/-^: n=15). \*\*\**P* < .001. ns, not significant.

### Tranexamic Acid Increases Fibrin Accumulation in WT but Not *CypD*^*-/-*^ or Antiplatelet Therapy-Treated Mice

As shown in Figure 3, fibrin accumulation was significantly increased in *Plg*^*-/-*^ mice. Consistent with this finding, we observed co-localization of recombinant plasminogen (rPlg) with fibrin immediately following laser injury in WT (Figure 6A) and *Plg*^*-/-*^ mice (Figure 6B). At later time points, rPlg staining was also observed away from the fibrin ring. Treatment of mice with TXA, an antifibrinolytic compound that inhibits Plg binding to fibrin (33), led to increased fibrin accumulation at sites of injury when compared to untreated controls (Figure 6C). In contrast to control mice, rPlg recruitment to hemostatic plugs in *CypD*^*-/-*^ mice was significantly reduced (Figures 6D,E), and TXA treatment only had a minimal effect on fibrin accumulation in *CypD*^*-/-*^ mice (Figure 6F).

**Figure 6.**
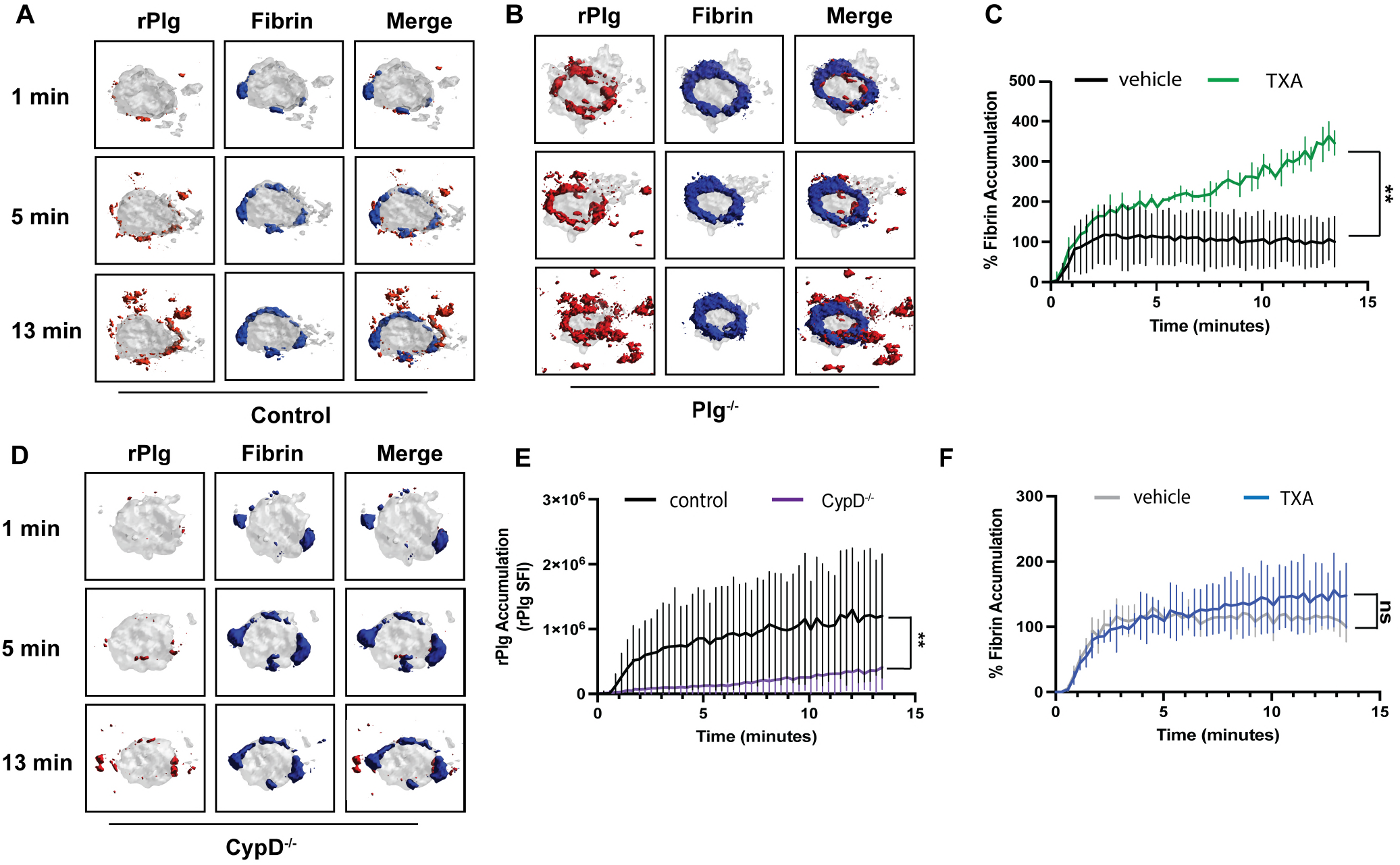
TXA does not increase fibrin accumulation in hemostatic plugs of *CypD*^*-/-*^ mice. Mice were pretreated with anti-fibrin-AF488 antibody, anti-GPIX-AF568 antibody, and AF647-conjugated, catalytically inactive recombinant plasminogen (rPlg). **(A-B)** Representative images (bottom/surface view) for fibrin (blue), platelet (grey) and plasminogen (red) staining at the indicated time points after laser injury in control **(A)** or *Plg*^*-/-*^ **(B)** mice. **(C)** Fibrin accumulation (as normalized to end point value in vehicle-treated mice) for vehicle or TXA-treated WT mice. (vehicle: n=18; TXA: n=16) **(D-F)** TXA effect on plasminogen recruitment and fibrin accumulation at sites of hemostatic plug formation in *CypD*^*-/-*^ mice. **(D)** Representative images (bottom/surface view) for fibrin (blue), platelet (grey) and plasminogen (red) staining at the indicated time points after laser injury. **(E)** Quantification of rPlg accumulation measured by rPlg (SFI ± SEM) for control and *CypD*^*-/-*^ mice. (control=9; *CypD*^*-/-*^ =10) **(F)** Fibrin accumulation (normalized as in D) for

TXA is clinically used to reduce bleeding, including in patients on antiplatelet therapy (34). We next studied the effect of aspirin and clopidogrel (dual antiplatelet therapy, DAPT) on fibrin accumulation in hemostatic plugs, both in the presence and absence of TXA. Platelets from DAPT-treated mice were markedly impaired in their ability to express PS after strong stimulation in vitro (Supplemental Figure 1). Fibrin accumulation at sites of injury was also significantly impaired in DAPT-treated mice (Figure 7A), and platelet density within the plug was reduced (Figure 7B). Similar to what we observed in *CypD*^*-/-*^ mice, TXA treatment did not improve fibrin accumulation in DAPT-treated mice (Figures 7C,D). Furthermore, TXA did not affect the bleeding time (Figure 7E) or blood loss volume (Figure 7F) following tail transection in clopidogrel-treated WT mice.

**Figure 7.**
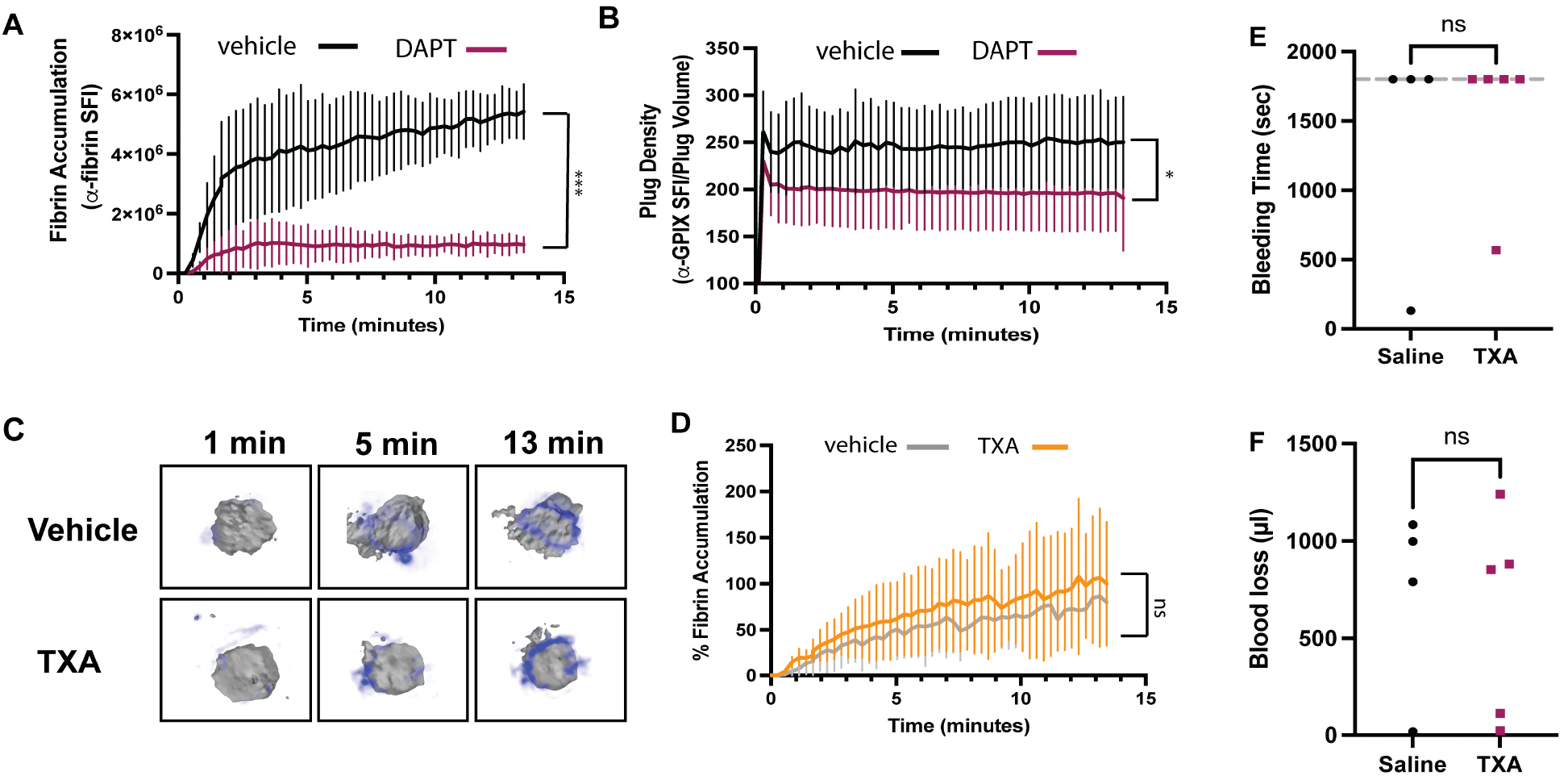
TXA does not improve hemostasis in antiplatelet drug-treated mice. Mice were pretreated with anti-fibrin-AF647 antibody and anti-GPIX-AF488 antibody. **(A)** Quantification of fibrin accumulation (SFI ± SEM) for vehicle and dual antiplatelet therapy (DAPT)-treated wild-type mice. (vehicle: n=18; DAPT: n=17) **(B)** Quantification of platelet plug density (SFI ± SEM) for vehicle and DAPT-treated wild-type mice (vehicle: n=12; DAPT: n=12) **(C)** Fibrin (blue, intensity view) and platelet (grey, surface view) accumulation at the indicated time points during hemostatic plug formation for DAPT-treated mice also receiving vehicle or TXA. **(D)** Fibrin accumulation (as normalized to end point value in vehicle-treated mice) for DAPT-treated mice also receiving vehicle or TXA. (vehicle: n=6; TXA: n=14). **(E, F)** WT mice were treated with clopidogrel and TXA (or vehicle control) and hemostasis was assessed by tail transection. **(E)** Bleeding time. (vehicle n=7, TXA n=16). **(F)** Blood loss volume (vehicle n=4, TXA n=5). \**P* < .05,\*\**P* < .01.

## DISCUSSION

Both platelets and ECs can provide a negatively charged, procoagulant surface required for assembly of coagulation complexes and subsequent thrombin generation and fibrin formation within hemostatic plugs. However, there is very limited information from intravital microscopy studies on the interplay between platelets and ECs with both the coagulation and fibrinolytic systems. We here described a novel 4D imaging platform to visualize and quantify hemostatic plug components in the saphenous vein of mice. Our studies suggest that platelet PS is more critical than EC PS for fibrin accumulation during hemostasis, that rapid onset fibrinolysis is one reason for limited fibrin accumulation on PS-expressing ECs, and that TXA has little effect on fibrin levels in hemostatic plugs of mice with impaired platelet PS expression.

We used mice deficient in CypD in all tissues and mice lacking CypD in platelets only to examine the relative contributions of platelet and EC PS during hemostatic plug formation. Our studies show that fibrin accumulation at a site of injury is predominantly observed at the plug-EC interface, consistent with what we and others previously showed with more conventional imaging techniques (26,35–37). Our 4D imaging method, however, allowed us to visualize both PS exposure and fibrin accumulation with improved spatiotemporal resolution. Specifically, we observed a first wave of PS exposure colocalizing with fibrin at the periphery of the platelet plug, right at the level of the endothelium. This first wave of PS exposure was not observed in mice lacking CypD in platelets only, indicating that platelets but not ECs play a crucial role in thrombin generation early during plug formation. Interestingly, subsequent PS expression, but not fibrin accumulation, was observed across the endothelium. These findings are partially consistent with those reported by Ivanciu et al. (17) and Schmaier et al. (13), who showed that prothrombinase complex and PS accumulation at sites of vascular injury is mostly found away from the platelet plug. However, these studies also showed that impaired platelet adhesion/ activation had little effect on fibrin accumulation at the injury site, suggesting that ECs play an unexpected role as a primary procoagulant surface during hemostatic and thrombotic plug formation. There are various potential explanations for these discrepant findings, including the type of injury, the length of the observation period following vascular damage, and the vessel itself. We generated penetrating lesions in the saphenous vein 50 μm in diameter that caused prolonged bleeding, consistent with a hemostasis model. In contrast, both Ivanciu et al. and Schmaier et al. created smaller, less defined injuries in cremaster arterioles that caused minimal RBC extravasation, even in the absence of functional platelets. Thus, our injuries are more severe and likely cause stronger platelet activation (including procoagulant activity) due to exposure to extracellular matrix and subendothelial TF. We also observed plug formation for a longer period of time following laser injury when compared to the studies by Ivanciu et al. and Schmaier et al. (13 minutes vs. less than 5 minutes). This is important as we observed a first wave of TF-dependent thrombin generation that was largely platelet-independent and lasted for about 2 minutes. Following this first wave was a second, platelet PS-dependent phase of fibrin formation. Ivanciu et al. also observed that fibrin accumulation plateaued at ∼3 minutes following laser injury in platelet-impaired, but not control mice. However, the reduction in fibrin levels did not reach significance, likely due to the short observation period. Schmaier et al. recorded for less than 3 minutes after laser injury, and thus likely would have missed a platelet PS-dependent phase of fibrin formation.

Our finding that PS-positive platelets play a predominant role in fibrin formation during hemostatic plug formation is in agreement with impaired hemostasis reported for mice deficient in TMEM16F in megakaryocytes and platelets only (14). In vitro, TMEM16F-deficient platelets showed a partial protection from agonist-induced PS exposure, leading to a significant reduction in thrombin generation (10,14). In vivo, mutant mice exhibited prolonged bleeding following tail transection. The observations described for megakaryocyte/platelet-specific TMEM16F knockout mice are similar to those reported for global TMEM16F knockout mice (11,38), suggesting that the mild bleeding phenotype observed in Scott syndrome patients is largely due to impaired procoagulant activity in platelets. So, what makes platelets special as a PS-positive surface that can support thrombin/fibrin formation? The relative level of PS exposure would be the simplest explanation for why platelets are a better procoagulant surface than ECs. Consistent with this hypothesis, aV fluorescence intensities were generally higher on the platelet surface when compared to surrounding ECs.

However, we also frequently observed PS hotspots, but not fibrin, away from the platelet plug. Thus, platelets likely provide more than just a PS surface to boost coagulation. It is well established that platelets also secrete coagulation proteins, such as FV and FXIII-A, as well as the fibrinolysis inhibitor, PAI-1 (19,39–41). Platelet FVa has more FXa cofactor activity than plasma-derived FVa (42), and hemostatic activity was demonstrated for both platelet and plasma FVa (43). Platelet FXIII-A is highly abundant in the cytoplasm and can be non-proteolytically activated by Ca^2+^ (44). Following cellular activation, released FXIII-A can reach local concentrations more than two orders of magnitude higher than the plasma FXIII concentration. FXIII-A is important for stabilizing fibrin by cross-linking individual fibers and by covalently attaching fibrinolysis inhibitors (45). Platelet PAI-1 has documented antifibrinolytic activity, both in vitro and in vivo (19). Aside from coagulation and fibrinolysis proteins, platelets also store and secrete very high levels of Zn^2+^, an essential co-factor of FXII. The Zn^2+^ concentration in platelet alpha granules is ∼25-60-fold higher than in plasma, and it was shown that platelet activation and granule release contribute to thrombin/fibrin generation and alter fibrin structure in vitro (46–48). Importantly, Zn^2+^ chelation in mice markedly impaired thrombus formation, similar to what was observed in *F12*^*-/-*^ mice (46). We also observed a marked reduction of fibrin accumulation in *F12*^*-/-*^ mice. Interestingly, fibrin accumulation was mostly observed at the interface between the platelet plug and the endothelium, i.e. the region where platelets encounter both thrombin and extracellular matrix components, and where platelets release their alpha granules. So, while both ECs and platelets bind FXII, platelets are in a unique position to activate FXII due to their ability to store and release large quantities of Zn^2+^. Further studies are required to delineate how these various mechanisms contribute to platelet-dependent fibrin accumulation within a hemostatic plug.

Our studies also provide critical new insights into the antagonistic balance between fibrin formation and breakdown during hemostatic plug formation. We observed significantly reduced fibrin accumulation in *F12*^*-/-*^ mice, a finding that seems surprising considering that patients and mice deficient in FXII do not bleed (49). However, there is considerable evidence that FXII is critical for thrombin/fibrin formation in atherothrombosis and venous thrombosis, i.e. situations where thrombus formation is initiated by small amounts of TF and propagated by the contact pathway (50–52). Our studies demonstrate a similar initiation/ propagation mechanism for hemostatic plugs, but they also indicate that the TF pathway is the primary activator of coagulation at sites of hemostasis. Consistently, FXII deficiency did not lead to bleeding, even though fibrin levels within plugs were significantly reduced. Another important finding of this work is that plasminogen is rapidly recruited to the hemostatic plug, mostly associated with the fibrin ring, resulting in conversion to plasmin and a rapid increase of fibrinolytic activity within the plug. This finding is consistent with previous work that showed rapid plasminogen recruitment to fibrin formed under flow conditions in vitro and in vivo (19,21,53). However, we also observed significant plasminogen binding in areas where we did not observe fibrin, likely due to other binding sites such as the recently described receptor, Plg-RKT (22), S100A10 (54), or PS-positive cells (19). Our studies with *CypD*^*-/-*^ mice showing reduced plasminogen recruitment to the plug area suggest a key role for PS in this process. However, we cannot exclude the possibility that rPlg recruitment to the vasculature, away from the fibrin ring, is mediated by trace amounts of fibrin(ogen) that are below the detection threshold of our system. TXA inhibits Plg binding to fibrin, thereby protecting fibrin from fibrinolysis during the hemostatic process. The increased spatiotemporal resolution of our model allowed us to visualize these processes in vivo, and we observed significantly increased fibrin levels in *Plg*^*-/-*^ mice or WT mice treated with TXA. Furthermore, we observed an increase in fibrin-positive area and fibrin ring thickness in these mice, suggesting that plasminogen action reduces fibrin accumulation within and on ECs directly adjacent to the platelet plug. TXA treatment did not significantly change fibrin accumulation in *CypD*^*-/-*^ mice or WT mice treated with antiplatelet drugs, however, consistent with the fact that fibrin accumulation was markedly impaired in these mice. These findings are of clinical relevance as TXA is routinely given to patients with bleeding complications, including those with inherited and acquired platelet function disorders. TXA has been shown to reduce bleeding and deaths due to postpartum hemorrhage (55), major trauma (56), and traumatic brain injury (57). However, no differences in World Health Organization (WHO) grade ≥2 bleeding, platelet transfusions or deaths were observed in patients with hypoproliferative thrombocytopenia (58). Furthermore, only mild positive effects of TXA treatment were observed in patients on APT, including reduced hematoma expansion in patients with brain hemorrhage (59) and reduced blood transfusions in patients undergoing coronary artery surgery (60). No measurable effects on death as an outcome were observed, and TXA treatment was associated with a higher rate of seizures. These studies led to questions on the balance of efficacy and safety of TXA in the prevention of bleeding in patients with defects in platelet count and/or function. Using two standardized models of hemostasis, we here show that TXA does not improve bleeding times in mice treated with the P2Y12 inhibitor clopidogrel, consistent with our observations that TXA does not lead to increased fibrin accumulation in situations where the platelet procoagulant response is significantly impaired, including APT.

In summary, we report on a novel 4D imaging platform to visualize and quantitatively assess hemostatic plug formation in mice. Utilizing this platform in combination with genetically modified mice, we demonstrate that platelets are the main procoagulant surface contributing to fibrin formation/accumulation within the plug. Our studies suggest that ECs immediately surrounding the injury area can also contribute to fibrin formation, but that this procoagulant activity is largely offset by rapid plasminogen recruitment and fibrinolysis. Importantly, TXA treatment was able to boost fibrin accumulation in control but not DAPT-treated mice, consistent with its limited pro-hemostatic activity reported for patients with platelet function disorders.

## Acknowledgements

The authors would like to thank Tanvi Rudran (University of North Carolina at Chapel Hill) for work in optimizing the tail bleed model, Dr. Peter Nigrovic (Brigham and Women’s Hospital) for use of cell lineage tracer mice, and Dr. Steven Grover for helpful discussions. This work was supported by National Institutes of Health, National Heart, Lung, and Blood Institute grants R35 HL144976 (W.B.), P01 HL151433 (W.B.), F31HL165935-01 (A.B.K), R35 HL155657 (N.M), R01 HL 160046 (M.J.F), and U01 HL14303 (M.J.F.).

## Authorship

Contribution: A.B.K designed research, performed research, analyzed data, and wrote manuscript; R.H.L designed research, performed research, analyzed data, and wrote manuscript; E.C.O analyzed data; S.R.J. performed research; P.Y.K, N.M. and M.J.F provided vital reagents or mice and provided valuable insights; D.S.P performed research; D.A. generated and provided data analysis software; and W.B designed research and wrote the manuscript.

## METHODS

### Materials

Anti-GPIX antibody (clone Xia.B4) was from Emfret Analytics. Anti-fibrin antibody (clone 59D8) was a kind gift by R. Camire (Children’s Hospital of Philadelphia). Annexin-V protein was generated and purified according to a protocol kindly provided by S. Krishnaswamy (Children’s Hospital of Philadelphia). Anti-human interleukin 4 receptor (IL4R) (clone 25463) antibody was from R & D Systems. Tranexamic acid was from Thermofisher. Aspirin and clopidogrel were from MedVet. Recombinant plasminogen was prepared as previously described (61–63).

### Animals

Wild-Type, hIL4Ra/GPIba-Tg (64), *TF*^*low*^ (28), *F12*^*-/-*^ (29), and *Plg*^*-/-*^ (30) mice have been described recently. mTmG x PF4-Cre mice (cell lineage tracing) were provided by Dr. Peter Nigrovic (Brigham and Women’s Hospital). Cyclophilin D (CypD)-/- mice were purchased from the Jackson Laboratories (JAX stock # 009071) (65). All mice were between 10-20 weeks in age. All strains were maintained on a C57BL6/J background in the mouse facility of the University of North Carolina at Chapel Hill. Where indicated, mice were treated with 300 mg/kg tranexamic acid (via intravenous injection) immediately prior to imaging or clopidogrel (5 or 25 mg/kg) and/or Aspirin (20mg/kg) via oral gavage 24 and 3 hours prior to imaging.

### Platelet Adoptive Transfer Model

Platelet adoptive transfer was performed as described previously (25). Briefly, endogenous platelets in hIL4Ra/ GPIba-Tg mice were depleted via retroorbital injection of anti-human IL4R antibody (2.5 μg/g body weight). After 2 hours, platelet depletion was confirmed by flow cytometry. Thrombocytopenic hIL4Ra/GPIba-Tg mice were transfused via retroorbital injection with washed platelets to reach a peripheral platelet count between 2.0 and 5.0 x108 platelets/ml.

### Saphenous Vein Laser Injury Model

Laser-induced injury to the saphenous vein was performed as described previously (26) with small modifications. Briefly, mice (10-20 weeks of age) were anesthetized by isoflurane inhalation (Veterinary Anasethesia Vaporizer, DRE Veterinary). The saphenous vein was exposed and the mouse was transferred to the imaging stage with a heating pad at 37°C. Relevant antibodies were injected (5 μg of each antibody) and a perfusion drip of saphenous vein buffer (physiological salt solution containing 132 mM NaCl, 4.7 mM KCl, 1.2 nM MgSO4, 2 mM CaCl2, and 18 mM NaHCO3 (pH 7.4) bubbled with 5% CO2/95% N2 for 10 minutes) was maintained at a rate of 200 ml/h. A perforating injury to the endothelium was induced via Ablate photoablation system (Ablate! photoablation system; Intelligent Imaging Innovations, Denver, CO, USA) with a 532nm pulse (70uJ; 100 Hz; duration of 10 ns with 7 ns interval between pulses), focused through a 20x water immersion objective lens (numerical aperture of 1). The resulting injuries were between 40 and 60 mm in diameter. Images were captured on a Zeiss Axio Examiner Z1 microscope equipped with a Yokogawa confocal scanning unit, CSU-W (Yokogawa Electric, Co), 488, 561, 640 nm lasers (Intelligent Imaging Innovations), and an Orca Flash 4.0 camera (Hamamatsu). Confocal images were acquired with SLIDEBOOK 6.0 software (Intelligent Imaging Innovations) using the following settings: 100ms exposure time, 4x4 binning, 7.5 μm Z-steps.

### Tail Clip Bleeding Model

Wild-type mice were treated with clopidogrel (5 mg/kg) by oral gavage 24 and 3 hours before the experiment. Mice were anesthetized with ketamine and xylazine (100 μg/g and 15 μg/g, respectively) followed by ketamine supplementation (500 μg) and placed on a heating pad maintained at 37° C. 5 minutes prior to tail clip, mice received intravenous injections of tranexamic acid (TXA) (300 mg/kg) or saline. Then, 2 mm of the distal tip of the tail was amputated using an ultrasharp cryostat blade and the tail was submerged in a 50 ml conical tube containing 45 ml of 37° C saline. Bleeding was monitored for 30 minutes, after which mice were euthanized. For determination of blood loss volumes, blood was pelleted and resuspended in 10 ml of filtered water to lyse RBCs, and hemoglobin concentration was measured at 416 nm using a microplate reader and compared against standards of known volumes of mouse whole blood.

### Image Analysis

Images were exported as TIFF-OME files from SlideBook 6.0 software. Image analysis was done with ImageTank software (Visual Data Tools, Inc). Accumulation of hemostatic plug components was quantified from sum fluorescence intensity (SFI) of relevant fluorescent channel. The density of hemostatic plug components was measured as SFI/ volume.

### Statistics

Statistical analysis was done using GraphPad Prism (GraphPad, Inc., LaJolla, CA, USA). All data is shown as mean +/- standard error of the mean. All analysis was done with welch t-test unless otherwise noted. P-values of less than 0.05 were considered significant. Only mice which did not rebleed were included in analysis.

### Study Approval

All experiments were approved by the Animal Care and Use Committee Tt of the University of North Carolina at Chapel Hill

