## Supplemental figures for "4D intravital imaging studies identify platelets as the predominant cellular procoagulant surface in a mouse model of hemostasis"

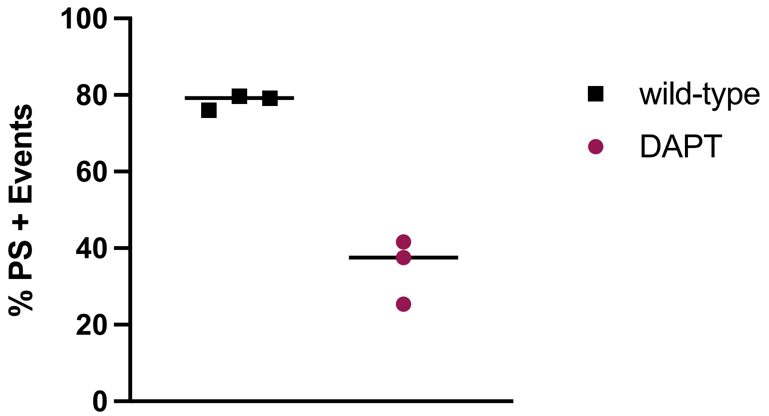


**Supplemental Figure 1. DAPT decreases PS exposure in vitro.**
Flow cytometric analysis of PS exposure measured by Annexin-V binding after activation with Par4P (250 μM) and Convulxin (100 ng/ml).
